# Head trauma impairs HPA-axis functions by increased R-loop structure and shortens telomeres

**DOI:** 10.1101/2024.05.29.596430

**Authors:** Zeynep Yılmaz Şükranlı, Serpil Taheri, Ecmel Mehmetbeyoğlu, Müge Gülcihan Önal, Mehmet Memiş, Begüm Er, Züleyha Karaca, Fatih Tanrıverdi, Kürsad Ünlühızarcı, Minoo Rassoulzadegan, Fahrettin Keleştimur

## Abstract

Traumatic brain injury (TBI) causes inflammation, one of the main causes of cellular aging. Telomere repeat-containing RNA (TERRA) hybridizes to telomere regions, forming R-loop structures and ensuring genome stabilization. Deregulation of R-loop homeostasis leads to genomic instability linked to neurodegenerative diseases and cancer. The hypothalamus-pituitary-adrenal (HPA) axis response is critical to maintaining homeostasis after TBI. We showed that the local increase in the transcription levels of the *Crh* and *Pomc* genes, in particular, suggests a defensive response through transcriptional alteration against mild TBI despite the decreased rate in the serum in the chronic phase. Additionally, changes in the transcription levels of TERRA and correlations with hormonal deficits after repetitive mTBI head trauma were observed. Telomere shortening and increased hybridized TERRA levels, especially after repeated mTBI in the chronic phase, suggest a possible disorder of genome stabilization and loss of cellular function in tissues of the hypothalamus, pituitary, and adrenal glands.

## INTRODUCTION

Traumatic brain injury (TBI) induces inflammation, resulting in elevated stress levels, a primary factor contributing to cellular aging (Sezgin Caglar *et al*, 2019; Taheri *et al*, 2022a, 2016; Greco *et al*, 2013). Despite numerous research, the mechanisms underlying hypopituitarism following head trauma remain unknown. Hypopituitarism has been shown to develop in boxers exposed to repetitive TBI. Pituitary dysfunction induced by TBI, whether repetitive or non-repetitive, is marked by partial or total impairment, and the most common hormonal defects are growth hormone (GH) and gonadotropin deficiencies (Tanriverdi *et al*, 2015, 2007; Kelestimur *et al*, 2004).

TBI has the potential to have a dual effect on the HPA axis. Cortisol secretion begins with increased inflammation by TBI and activation of the Hypothalamus-Pituitary-Adrenal (HPA) axis. In the long term, pituitary insufficiency is one of the TBI-related diseases, and it is defined as the disturbance of the secretion of one or more hormones secreted by the pituitary gland after TBI. In our recent study (Taheri *et al*, 2022a), we found increased apoptosis markers in the adrenal glands during the active phase and in the pituitary during the chronic phase of head trauma. This may indicate the loss of the secretory function in the cell. In addition, increased cortisol levels and inflammation are implicated in pathologies associated with telomere alteration (Kuzminskaite *et al*, 2021; Bazaz *et al*, 2021). The disruption of regulation and shortening of telomeres leads to loss of cellular function or death. However, we do not yet know the effects of the associated mechanisms of mTBI and r-mTBI on telomer regulations.

Telomeres are nucleoprotein structures located at the ends of chromosomes that preserve the stability of the genome. We know that telomeres shorten with age and excessive stress. They are structurally and functionally different from other chromosomal DNA sequences. These are the simple heterochromatic structures associated with the Shelterin complex, composed of six distinct proteins and TTAGGG repeat units. Telomere length (TL) is maintained by telomerase. Telomeres shorten with each somatic cell division, and when the cell crosses the hay boundary, the shortening ceases, causing growth arrest and aging. According to the literature, an increase in cortisol triggers both telomere shortening and aging (Lipps & Rhodes, 2009).

For many years, telomeres were considered transcriptionally silent. Since the discovery of non-coding RNAs of telomeric regions, these long non-coding RNAs, Telomeric Repeat Containing RNA (TERRA), have been assigned a role in protecting both the ends of chromosomes from degradation and a role ensuring the stabilization of the chromosomes (De Rosa & Opresko, 2023; Mehmetbeyoglu *et al*, 2022).

The R-loops, called RNA: DNA hybrids, form naturally or due to replication stress or DNA damage. R-loops are needed to be removed appropriately for genomic stability. R-loop homeostasis disruption, often observed in neurodegenerative diseases and cancer, leads to genomic instability. TERRAs play a vital role in this process by promoting homologous recombination in shorter telomeres, triggering a DNA damage response. Short telomeres accumulate TERRA R-loops regardless of cell cycle regulation, preserving telomere length and facilitating homologous recombination in response to damage signals. In long telomeres, eliminating TERRA R-loop clusters involves the telomeric protein Rif2 capturing ribonuclease HII (RNase HII). Conversely, in short telomeres, RNase HII is not eliminated and degrades TERRA R-loops. Telomeric proteins suppress TERRA R-loop aggregation, with RNase HII levels higher in long telomeres, decreasing as telomeres shorten—a critical feedback mechanism. As a result, the levels of transcribed and rehybridized TERRA from short telomeres may differ from those in long telomeres (Rassoulzadegan *et al*, 2021; Feretzaki *et al*, 2020; Graf *et al*, 2017). It is essential to examine the roles of TERRAs in diseases.

In our previous studies, we found that TBI induces epigenetic changes in acute and chronic phases and that these epigenetic changes last in some chronic patients (Taheri *et al*, 2022a, 2016). Long noncoding RNAs (lncRNAs) are one of the epigenetic mechanisms. TERRA is a lncRNA that plays a crucial role in TL and genome integrity (Isken & Maquat, 2009).

Here, the question was whether TBI affected telomere length or telomeric RNA (TERRA) expression in tissues composing the HPA axis. This study created a mouse model of single mild (mTBI) and repetitive mild TBI (r-mTBI). Next, we examined R-loop formation and telomere regulation in the hypothalamus, pituitary, and adrenal glands over transcripts and hormonal responses related to HPA, HPG, and IGF-1-GH axes in acute and chronic phases after TBI.

## MATERIALS AND METHODS

For this study, the mTBI and r-mTBI model was created in 2-month-old adult Balb/c mice (5 males, 5 females in each group, a total of 20 males, 20 females) with the approval of the Ethics Committee of Animal Experiments of Erciyes University (decision number: 19/127). The mice were sacrificed on the specified days after mTBI, and samples of hypothalamus, pituitary, and adrenal tissues and blood were collected. DNA and RNA samples were isolated from the hypothalamus, pituitary gland, and adrenal glands. DNA: RNA hybrids were isolated from DNA samples. TL was determined in DNA samples. Next, expression levels of free TERRA (fTERRA) were measured in total RNA samples, and hTERRA expression levels were determined in DNA: RNA hybrids isolated from DNA samples. The expression levels of *Corticotrophin Releasing hormone (Crh), Proopiomelanocortin (Pomc), Corticosterone (Cort), Gonadotropin Releasing Hormone (Gnrh), Growth Hormone (Gh), Insulin-Like Growth Factor-1(Igf1)* and *Arginine Vasopressin (Avp)* genes were determined in the hypothalamus, pituitary, and adrenal glands. Hormone levels were determined by ELISA d in serum samples. (The experimental flowchart can be seen in Figure 1). Our study design ensured a homogeneous group of animals with consistent housing conditions and minimized the impact of extraneous variables. The animals were cared for and treated according to the Principles of Laboratory Animal Care (European rules). Female mice were housed together from birth to minimize hormonal fluctuations (Marchlewska-Koj *et al*, 1983; Taheri *et al*, 2022b; Dalal & Khanna-Chopra, 2001).

**Figure 1.**
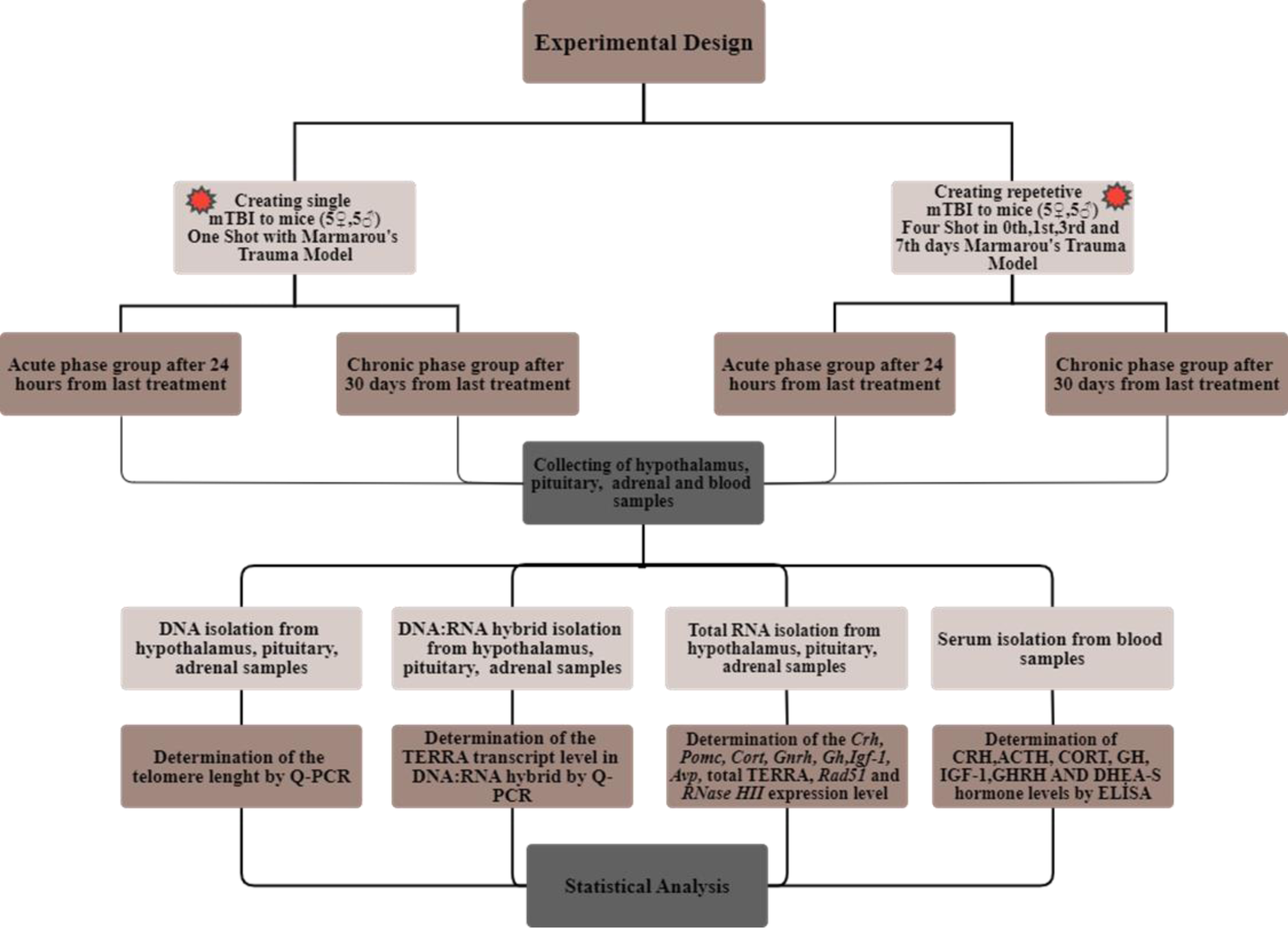
Experimental design.

### Creating the TBI Model in Mice

The TBI was applied in 2-month-old *Balb/c* mice according to the trauma model of Marmarau et al. (Marmarou *et al*, 1994; Tweedie *et al*, 2013). After anesthesia, a surgical incision was made to the scalp of the mouse with a sterile scalpel to expose the skull for the trauma procedure. To apply mild traumatic brain injury (mTBI), the mouse was placed under the Marmarau trauma device. This installation consists of an 80 cm pipe and an area under which the mouse is placed. A plastic disc was placed on the surgically exposed skull of the anesthetized mouse, and the pipe was lowered to fit perpendicularly on top of the plastic disc. Later, a 30-gram weight was dropped from the top of the pipe. This process was performed once to generate mTBI (Marmarou *et al*, 1994; Tweedie *et al*, 2013).

While establishing a r-mTBI model, mTBI was applied by firing a total of 4 times on days 0, 1, 3, and 7, including the first head trauma, for both acute and chronic groups. After the trauma, the scalp incisions were sutured with 26 mm and 75 mm sutures (Surgisorb, England), and the mice were placed back into their cages. After the trauma model was established, vital functions were monitored. The acute r-mTBI and acute m-TBI groups were sacrificed 24 hours after the last application, and the chronic r-mTBI and chronic mTBI groups were sacrificed 30 days after the last application. The mice designated for the control group were not exposed to TBI and were sacrificed at 3 months old. Hypothalamus, pituitary, and adrenal tissues were collected from sacrificed animals to isolate DNA, DNA: RNA hybrid, and total RNA. All results are derived from extracts from the whole hypothalamus, pituitary glands, and adrenal glands. Samples were stored in 1.5mL Eppendorf® Safe-Lock microcentrifuge tubes (Eppendorf, USA, CT, Catalog No. 0030123611) at − 80° C in Purezol until use. Blood samples were collected from mouse hearts to determine hormone levels, and serum isolations were performed (The experimental flowchart can be seen in Figure 1).

### Total RNA Isolation

Half of the collected tissues were used for RNA isolation. Tissues were collected in 500 µl of Purezol (Biorad, USA, CA, Cat No: 7326890) and homogenized with a 2 ml syringe. Subsequently, total RNA isolation was carried out according to our previous study and the manufacturer’s instructions (Taheri *et al*, 2022b; Tufan *et al*, 2023).

### DNA Isolation

The other half of the tissues was used to obtain DNA and DNA: RNA hybrid (TERRA). Each tissue was placed in 600 µl of Cell Lysis Buffer containing 20 mM Tris (pH 8), 0,5% SDS, 0.1 M EDTA, and Proteinase K 450µg/ml and carefully vortexed, then allowed to incubate at 56°C overnight. The next day, two Eppendorf tubes were prepared for each tissue, and 300 µl was dispensed into each Eppendorf tube from the tissue we incubated overnight, one tube for DNA isolation and one tube for DNA: RNA hybrid isolation. DNA isolations were performed according to the ammonium acetate method (Mehmetbeyoglu *et al*, 2022).

### DNA: RNA Hybrid Isolation

During the DNA isolation step, samples intended for DNA:RNA hybrid isolation were dried and nuclease-free water was not added. Instead, 75 µl DNA Digestion Buffer and 5 µl DNAase (Zymo Research, USA, CA, Cat No: E1010) were added to the dried pellet, vortexed for 10-15 seconds, and incubated at room temperature for 15 minutes. After incubation, 300 µl of Purezol (Biorad, USA, CA, Cat No: 7326890) and 180 µl of chloroform were added and isolation procedures were performed according to the manufacturer’s instructions. In this way, DNA was removed through DNase treatment, resulting in the isolation of DNA-associated RNAs (DNA:RNA hybrids) (Mehmetbeyoglu *et al*, 2022).

### cDNA Synthesis

Total RNA and DNA:RNA hybrid samples were reverse transcribed into complementary DNA (cDNA) using Evoscript universal cDNA Master Kit (Roche, Germany, Mannheim, Cat No: 07912439001) in final reaction volumes of 20 µL. All reactions were performed as specified in the manufacturer’s protocol. The cDNA samples were stored at −80°C until they were analyzed by Q-PCR (Mehmetbeyoglu *et al*, 2022).

### Q-PCR

Quantitative real-time PCR (q-PCR) reactions were performed using the Light Cycler 480 II high-throughput Real-Time PCR system (Roche, Germany, Mannheim). The cDNAs were diluted with nuclease-free water at a 1:5 ratio. Syber Green Master (Roche, Germany, Mannheim, Cat No: 04707516001) was used to determine the transcript levels of *Crh, Pomc, Cort, Gnrh, Gh, Igf1* and *Avp* genes and TERRA expression levels in total RNA and DNA: RNA hybrids.

The reaction mix was prepared according to the manufacturer’s instructions. Mouse beta-actin (ACTB) was used as a reference gene. Changes in gene expression were determined using the 2^-ΔΔCt^ relative quantification method in all groups. The primers used in the study are listed in Supplemental Table S1 (Mehmetbeyoglu *et al*, 2022).

### Telomere Length (TL) Measuring

To determine the telomere lengths (TL) of DNA isolated from tissues, telomere standard, and 36B4 standard were calculated via Real-Time PCR method using the SYBR Green I Master mix (Roche, Germany), and telomere and 36B4 primers, provided in Supplemental Table 1. Two separate mixtures for the number of samples and standards were prepared in separate tubes for telomeres and 36B4 primers. The reaction mix was prepared according to our previous study and the manufacturer’s instructions. Six distinct sets of standards were created by diluting the telomere and 36B4 standards. 1µl PGL3 Basic Vector (Promega, USA, Cat No: E1751) plasmid was added into all prepared standards. Once the reaction mix was distributed into the wells, 4 µl of the prepared standards were added sequentially. Thermal cycling conditions were set according to the manufacturer’s protocol The standard curve was plotted with the Ct values obtained for the telomere and 36B4 standards. Using the method suggested by (Mehmetbeyoglu *et al*, 2022; O’Callaghan & Fenech, 2011), the TL was calculated with the Ct values of each tissue.

### ELISA Assay

Blood samples collected from mice were centrifuged at 3000 rpm for 20 minutes for phase separation, and serum samples from the upper phase were transferred to the new Eppendorf tubes and stored at −80°C until the start of the study. Serum levels of hormones were studied using CRH (Ylbiont, Shanghai, Cat No: YLA0443MO), GH (Sunred, Shanghai Cat No: 201020677), ACTH (Ylbiont, Shanghai, Cat No: YLA1790MO), GHRH (Ylbiont, Shanghai, Cat No: YLA0164MO), IGF-1(Sunred, Shanghai, Cat No:201020038), CORT (Ylbiont, Shanghai, Cat No: YLA0342MO), DHEA-S (Ylbiont, Shanghai, Cat No: YLA0536MO) and Estrogen (Ylbiont, Shanghai, Cat No: YLA0030MO) ELISA kits. Serum hormone levels were determined according to our previous study and the manufacturer’s instructions (Akkar *et al*, 2022).

### Statistical Analysis

Once the results were obtained, comparisons between the experimental groups were made. Data conformity to normal distribution was assessed by histogram, q-q plots, and Shapiro-Wilk test. Correlation analyses were performed with One Way ANOVA, Kruskal Walls, student t-test, Mann Whitney-U test, and Pearson and Spearman tests, depending on whether the data showed a normal distribution or not. Data were analyzed using SPSS software version 22 (IBM, USA) and Graph-Pad Prism 8. Results with p values <0.05 were considered statistically significant (Mehmetbeyoglu *et al*, 2022).

## RESULTS

### Following TBI, Telomere length (TL) shortens and the amount of TERRA hybridized with DNA increases in the hypothalamic-pituitary-adrenal axis

Telomere lengths (TL) in the hypothalamus, pituitary gland, and adrenal glands were compared between groups after TBI in the acute and chronic phases. Figures 2 A, 2B, 2C show that TL is significantly shortened in the mTBI and r-mTBI groups compared with the control group in the hypothalamus, pituitary, and adrenals (Supplemental Table S2).

**Figure 2.**
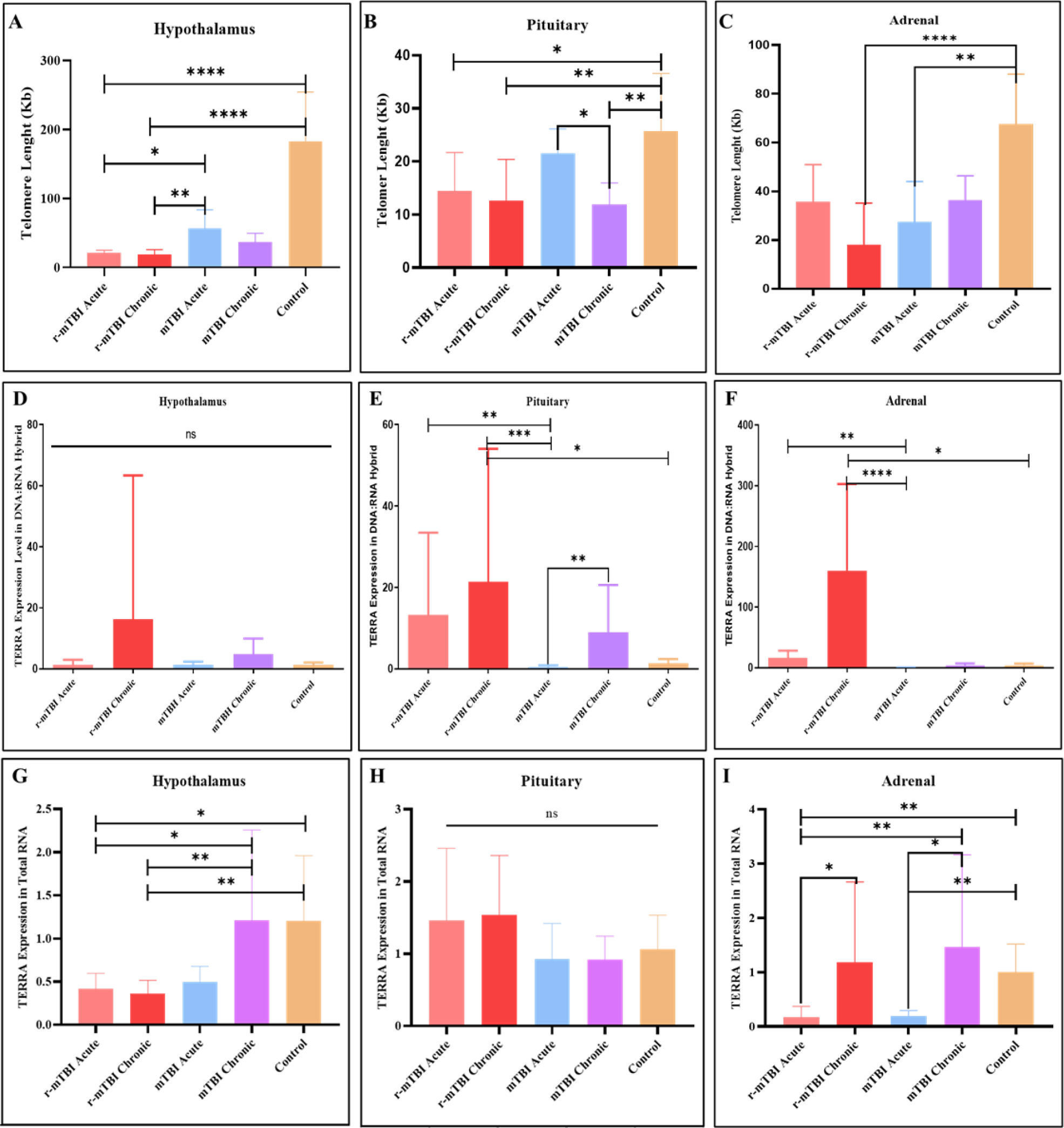
Differences in telomere length (TL), hTERRA, and fTERRA levels between groups in the hypothalamus, pituitary, and adrenal tissues (**ns**= p≥0.05, **\***p<0.05, **\*\***p<0.01 **\*\*\*** p<0.001 **\*\*\*\***p≤0.0001).

Interestingly, DNA-associated TERRA (hTERRA) levels in the hypothalamus, pituitary, and adrenal glands after mTBI increased the most in the r-mTBI group compared to the other TBI groups and the control group (Figures 2D, 2E, 2F, Supplemental Table S2).

In the hypothalamus, free TERRA transcript levels (fTERRA) decreased in the mTBI-acute, r-mTBI acute and r-mTBI chronic groups compared to the control group. However, there was no significant difference between the mTBI chronic and control groups. While there was no significant difference found in transcript levels in the pituitary, fTERRA transcript levels in the adrenal decreased in the acute mTBI and r-mTBI groups compared to the control group, yet increased in the mTBI chronic and r-mTBI-chronic groups (Figure 2G, 2H, 2I, Supplemental Table S2).

### *Rad51* and *RNaseHII* are altered after TBI: regulators of telomere length and TERRA profiles

Changes in TL and TERRA transcript levels, especially in the hybrid fraction attached to DNA, suggest a need to examine regulatory factors. Two important players in regulating TERRA binding to DNA and TL are *Rad51* and *RNase HII* gene products.

When *Rad51* transcript levels were compared between groups; there was no difference in the hypothalamus, but there was an increase in the r-mTBI-chronic group in the pituitary. Additionally, the decrease in acute groups (mTBI-acute and r-mTBI-acute) in the adrenal glands was significant (Figure 3A, 3B, 3C).

**Figure 3.**
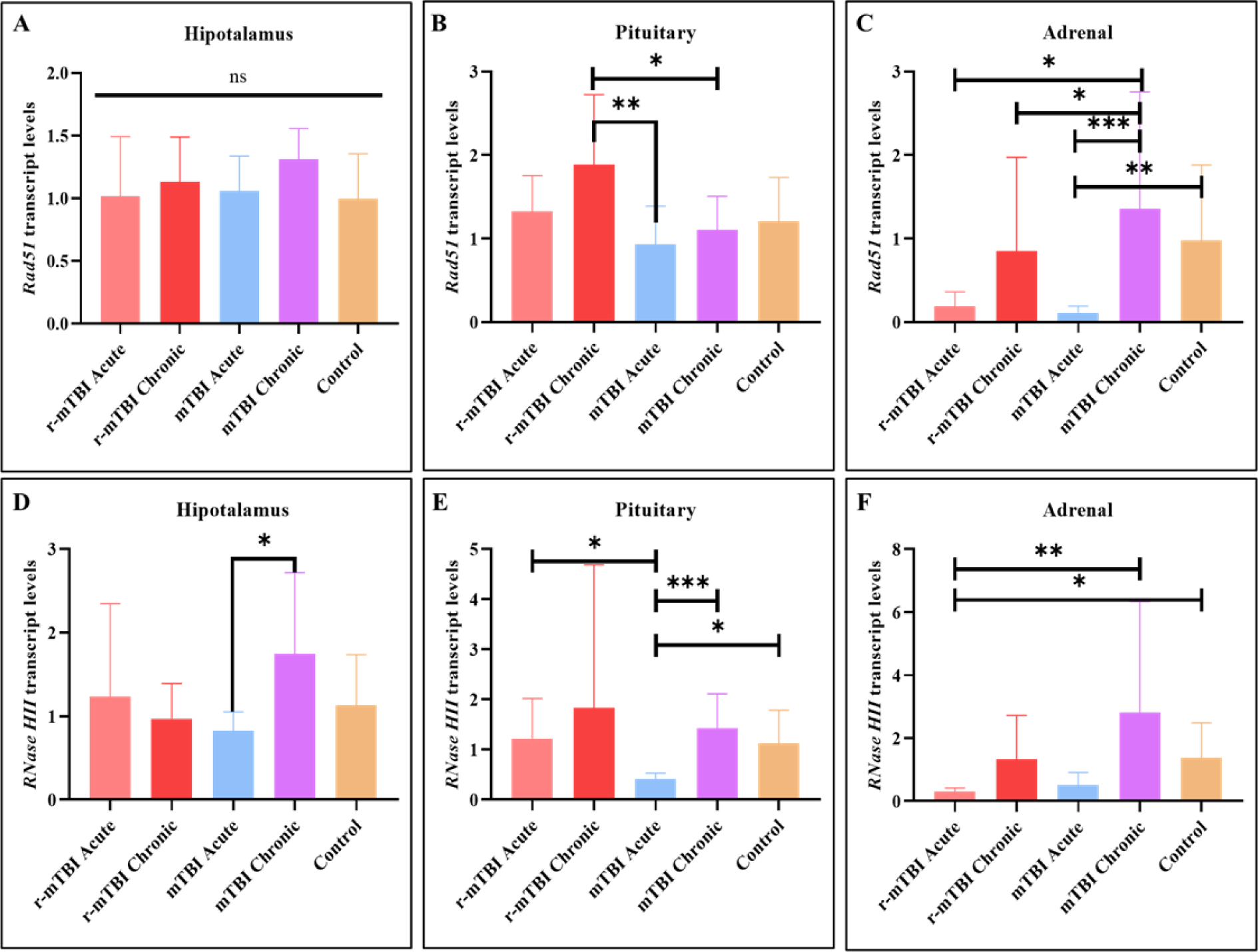
Differences in transcription levels of *Rad51* and *RNase* HII genes in the hypothalamus, pituitary, and adrenal glands between groups (**ns**= p≥0.05, **\***p<0.05, **\*\***p<0.01 **\*\*\*** p<0.001 **\*\*\*\***p≤0.0001).

When *RNaseHII* transcript levels were compared between groups; in the hypothalamus, the mTBI-acute group decreased according to the m-TBI-chronic group, and in the pituitary, it decreased in the mTBI-acute group compared to almost all groups. The decrease in *RNaseHII* transcript levels in the adrenals in the acute groups was significant (Figure 3D, 3E, 3F, Supplemental Table S3).

Graphical Figure 4 shows the TL and hTERRA, fTERRA, *RNaseHII*, *Rad51* transcript level fold changes in the hypothalamus, pituitary, and adrenal glands between the trauma groups after mTBI and r-mTBI compared to the control group. Increases, decreases and correlations are shown in the hypothalamus, pituitary, and adrenals in Figure 4, with fold change values for the control group considered zero. When the TL is compared in the hypothalamus, pituitary, and adrenals in the acute r-mTBI and r-mTBI-chronic groups, the TL shortening was most prominent in the hypothalamus among all groups (Figure 4A). hTERRA levels were highest in the r-mTBI-chronic group in all tissues compared to the other experimental groups (Figure 4A, 4B, 4C).

**Figure 4.**
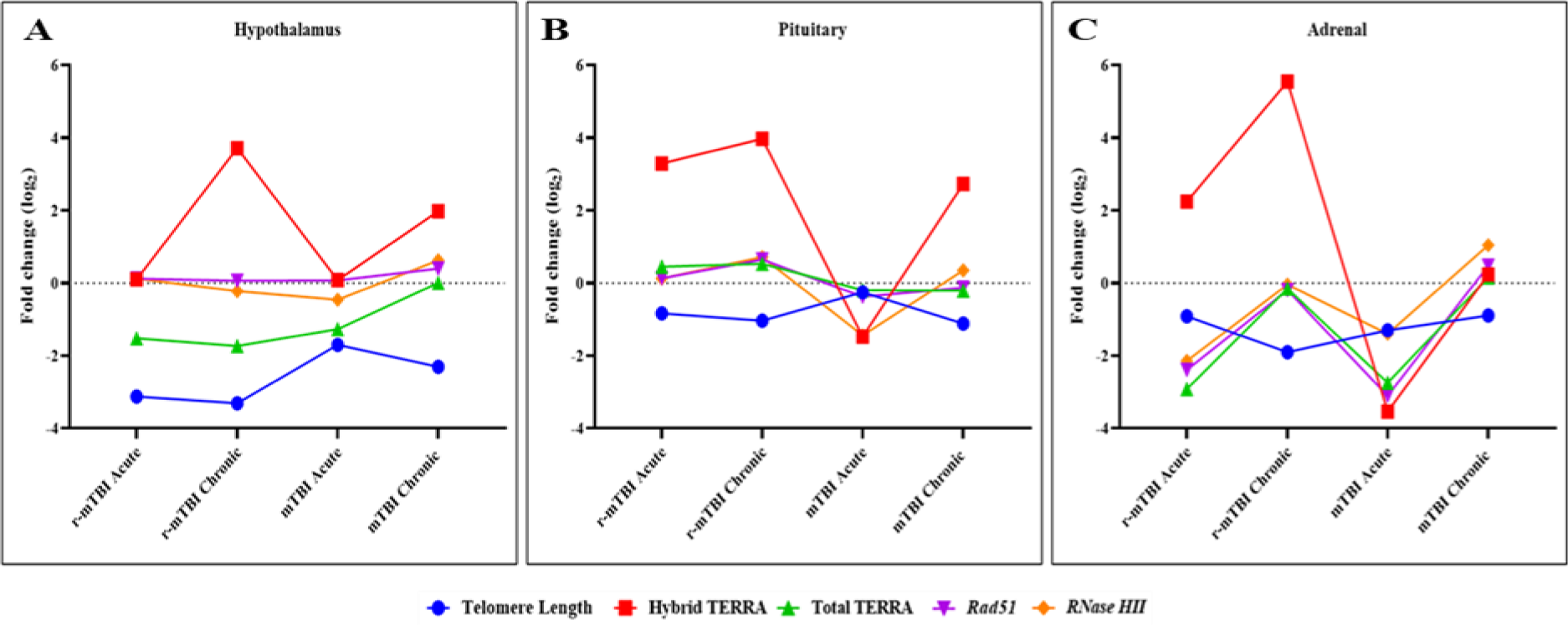
Graphical representation of telomere length (TL), hTERRA, fTERRA, *RNaseHII*, and *Rad 51* transcript levels in the hypothalamus, pituitary, and adrenal between groups after mTBI and r-mTBI. **A.** Telomere length (TL), hTERRA, fTERRA, *RNaseHII*, and *Rad 51* transcript levels in the hypothalamus. B. Telomere length (TL), hTERRA, fTERRA, *RNaseHII*, and *Rad 51* transcript levels in the pituitary. C. Telomere length (TL), hTERRA, fTERRA, *RNaseHII*, and *Rad 51* transcript levels in the adrenals.

The correlation between fTERRA, *Rad51*, and *RNase HII* transcript levels in adrenals after TBI is quite remarkable. When comparing the levels of hTERRA among different tissues, the most significant increase is observed in the adrenals during the chronic phase following repetitive mild traumatic brain injury (r-mTBI) (Figure 4C).

### Differences in HPA axis-related transcription and hormone levels in the Hypothalamus, Pituitary, and Adrenal glands after mTBI and r-mTBI

Figure 5 shows the transcription levels of HPA axis-related genes in the hypothalamus, pituitary and adrenal after mTBI. *Crh* transcript levels increased 150-fold in the r-mTBI-chronic group compared to the control group in the hypothalamus (Figure 5A). In the pituitary, *Crh* transcript levels increased in the r-mTBI-acute group compared to all other groups (Figure 5B). Although *Pomc* transcript levels did not differ significantly between groups in the pituitary, they were increased in the r-mTBI-chronic group in the hypothalamus and in the mTBI and r-mTBI chronic groups in the adrenals compared to the control and acute groups (Figure 5D, 5E, 5F). In the adrenals, it was notable that while the transcript levels of *Crh, Pomc, Avp*, and *Cort* increased in the chronic groups compared to the control group, they decreased in the acute groups (Figure 5F, 5I, 5L, Supplemental Table S4).

**Figure 5:**
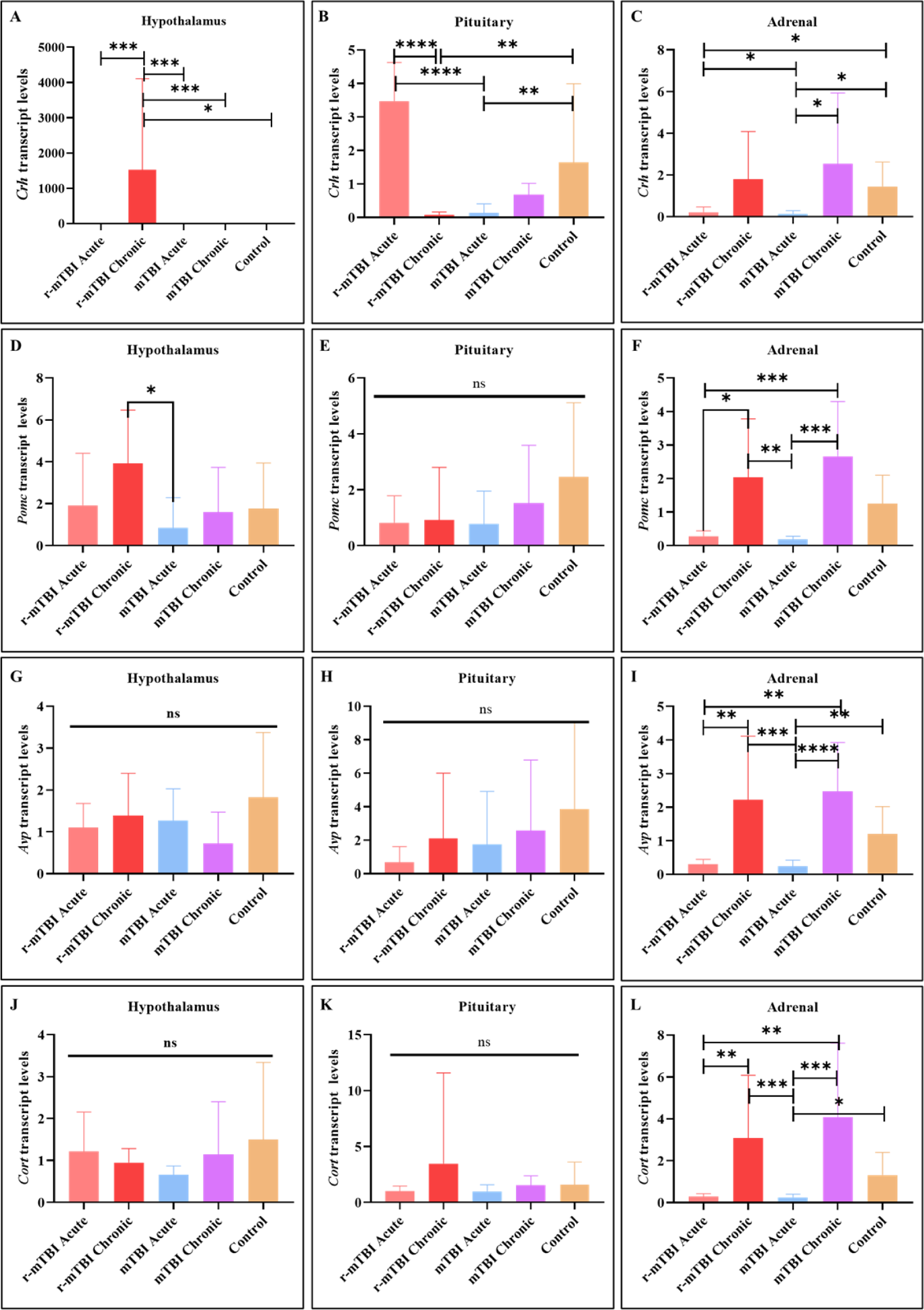
Differences in transcription levels of *Crh, Pomc, Avp* and *Cort* genes in hypothalamus, pituitary and adrenal tissues after mTBI and r-mTBI (**ns**= p≥0.05, **\***p<0.05, **\*\***p<0.01 **\*\*\*** p<0.001 **\*\*\*\***p≤0.0001).

When serum levels of CRH, ACTH, and Corticosteron (CORT) hormones were compared between groups after mTBI and r-mTBI; CRH levels decreased in the r-mTBI-chronic group compared to the control group (Figure 6A and Supplemental Table S5), while ACTH levels increased in the r-mTBI-acute group compared to the r-mTBI-chronic group (Figure 6B and Supplemental Table S5). CORT hormone levels increased in the r-mTBI-acute group compared to other groups (Figure 6C and Supplemental Table S5).

**Figure 6.**
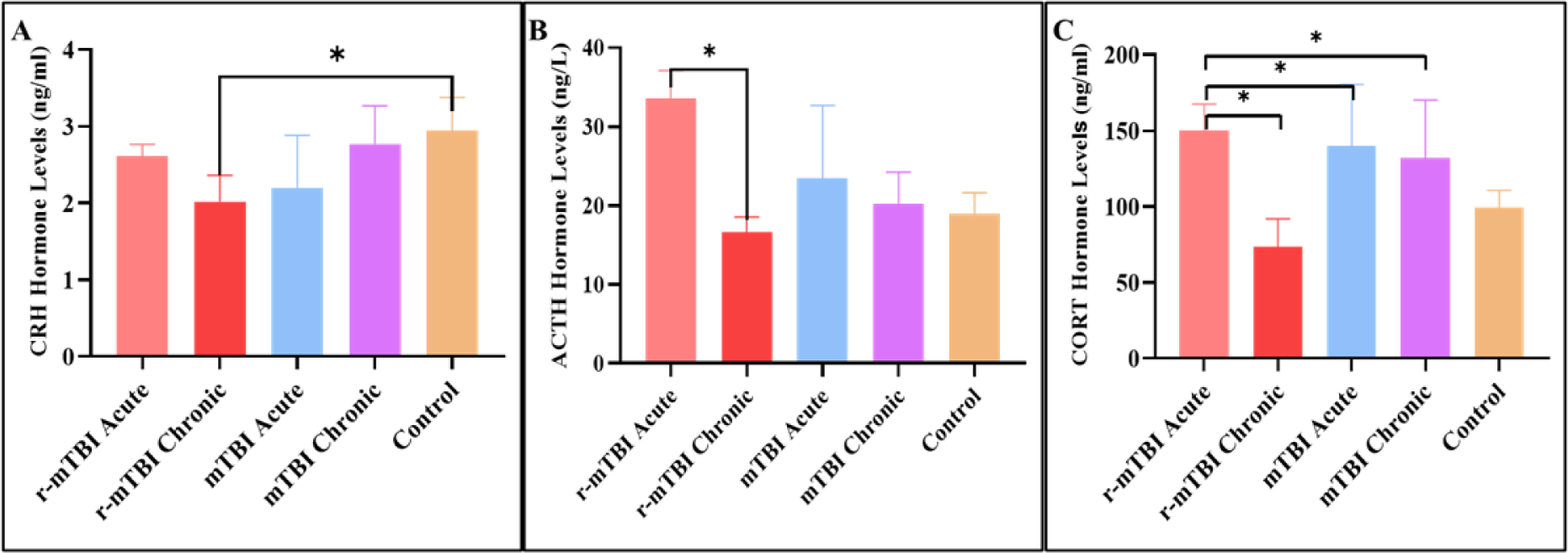
Differences in CRH, ACTH, and CORT hormone levels between the groups in serum after mTBI and r-mTBI (**ns**= p≥0.05, **\***p<0.05, **\*\***p<0.01 **\*\*\*** p<0.001 **\*\*\*\***p≤0.0001; **A:** Acute, **C:** Chronic)

### Differences in HPG axis-related transcripts in the Hypothalamus, Pituitary, and Adrenal glands and DHEA-S serum hormone levels after mTBI and r-mTBI

After mTBI and r-mTBI, *Gnrh* transcript levels in the hypothalamus, pituitary, and adrenal glands were compared between groups. In almost all hypothalamus groups, transcript levels decreased compared to the control group, but the most significant decrease was recorded in the mTBI-chronic group. Although there was no difference between groups in the pituitary, *GnRH* transcript levels in the mTBI-chronic and r-mTBI-chronic groups increased compared to the control group in the adrenal glands (Figure 7A, 7B, 7C, Supplemental Table S6).

**Figure 7.**
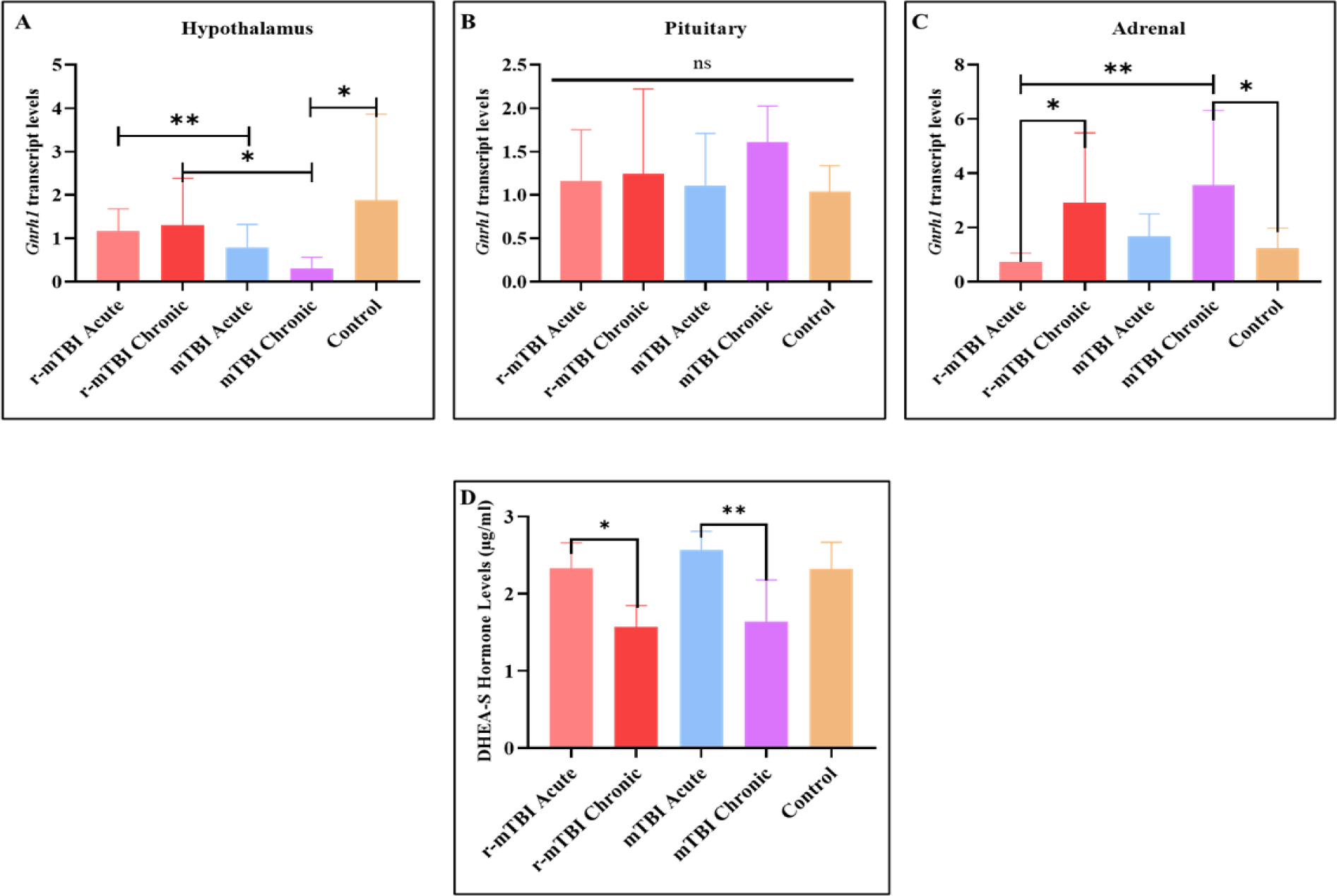
Differences in hypothalamus, pituitary, and adrenals *GnRH* transcription levels in the and DHEA-S hormone levels after mTBI and r-mTBI (**ns**= p≥0.05, **\***p<0.05, **\*\***p<0.01 **\*\*\*** p<0.001 **\*\*\*\***p≤0.0001; **A:** Acute, **C:** Chronic).

When serum DHEA-S hormone levels were compared between groups, it was determined that the mTBI-chronic and r-mTBI-chronic groups decreased compared to the acute groups (Figure 7D).

### Differences in GH-IGF-1 axis-related transcripts in hypothalamus, pituitary, adrenals, and serum hormone levels after mTBI and r-mTBI

*Gh* and *Igf-1* transcript levels were compared in the hypothalamus, pituitary, and adrenal glands between the mTBI and r-mTBI groups. While *Gh* transcript levels in the hypothalamus decreased in the r-mTBI-acute and r-mTBI-chronic groups, no significant differences were found in the pituitary. However, Gh transcript levels increased in the chronic groups compared to the control group, while they decreased in the acute groups in the adrenals after mTBI head injury (Figure 8A, 8B, 8C, Supplemental Table S7).

**Figure 8:**
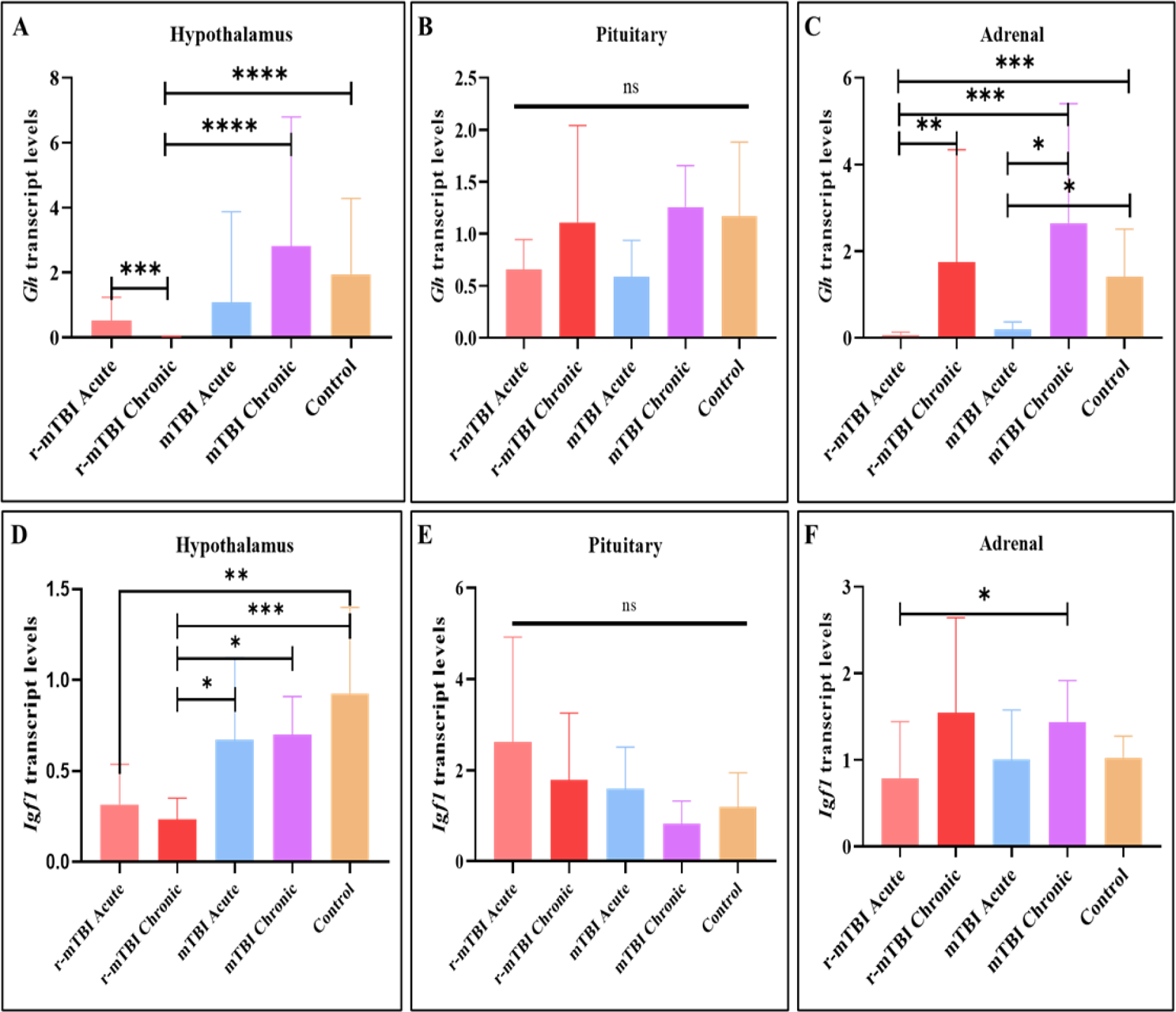
Differences in *Gh* and *Igf-1* transcript levels in the hypothalamus, pituitary and adrenal gland after mTBI and r-mTBI (**ns**= p≥0.05, **\***p<0.05, **\*\***p<0.01 **\*\*\*** p<0.001 **\*\*\*\***p≤0.0001).

*Igf-1* transcript levels decreased in the r-mTBI-acute and r-mTBI-chronic groups in the hypothalamus after TBI, and there was no difference between the pituitary and adrenal groups (Figure 8A, 8B, 8C).

Finally, hormone levels related to the GH-IGF-1 axis-were compared between groups after mTBI; there was no difference between serum GH and IGF-1 levels, but serum GHRH levels were found to be increased in the r-mTBI-acute group compared to other groups (Figure 9A, 9B, 9C and Supplemental Table S8).

**Figure 9.**
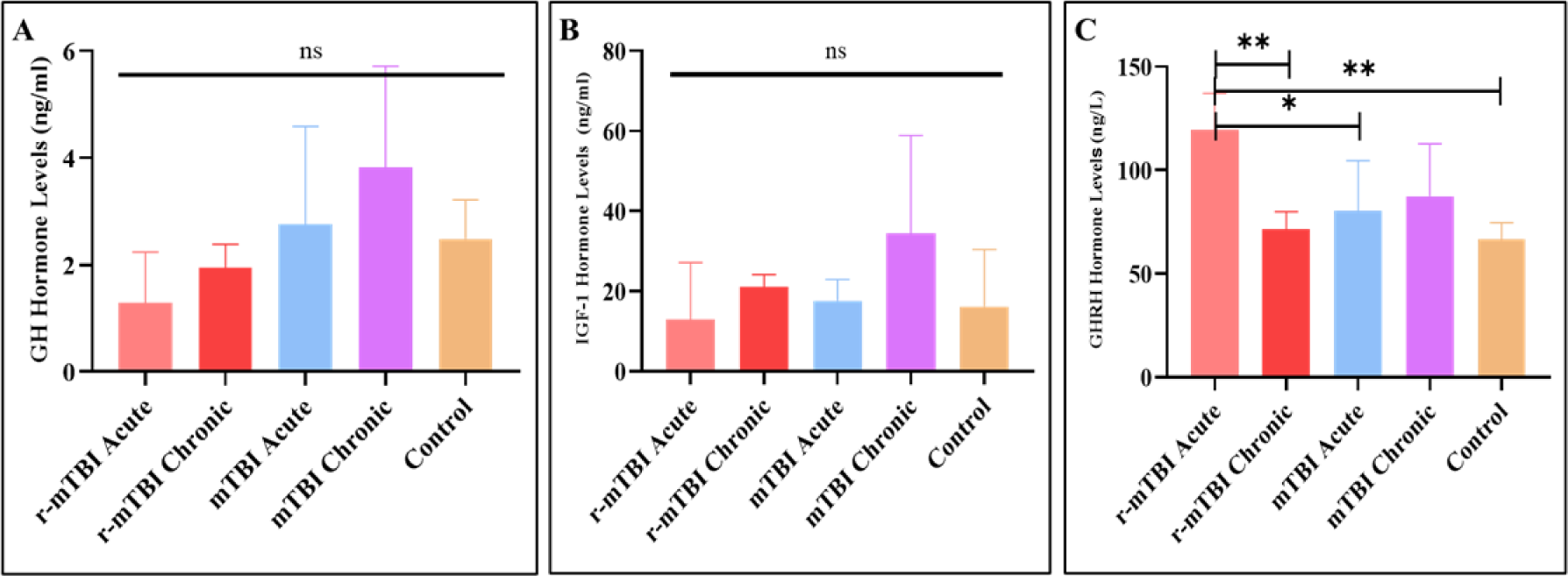
Differences in serum GH, IGF-1 and GHRH levels between groups after mTBI and r-mTBI (**ns**= p≥0.05, **\***p<0.05, **\*\***p<0.01 **\*\*\*** p<0.001 **\*\*\*\***p≤0.0001).

Finally, hormone levels related to the GH-IGF-1 axis-were compared between groups after mTBI; there was no difference between serum GH and IGF-1 levels, but serum GHRH levels were found to be increased in the r-TBI acute group compared to other groups (Figure 9A, 9B, 9C).

Correlations between telomere length, fTERRA, and hTERRA levels, and the levels of hormones released from the hypothalamus, pituitary, and adrenal glands of mTBI mouse groups. When hypothalamic telomere length, fTERRA, and hTERRA levels were compared with CRH and GHRH hormone levels, r-mTBI chronic group had the most shortening of TL and the highest increase hTERRA levels in the hypothalamus, and lowest serum concentrations of CRH and GHRH compared to other groups (Figure 10A). When telomere length, fTERRA and hTERRA levels in the pituitary are compared with GH and ACTH levels; While hTERRA was at its lowest level in the mTBI acute group, ACTH and GH levels were found to be close to the control group. In r-mTBI chronic group demonstrated the shortest telomere length and highest hTERRA levels, GH and ACTH serum levels were decreased (Figure 10B). In the adrenal gland, r-mTBI chronic group had the highest hTERRA levels and, CORT and DHEAS levels decreased, and telomere length shortened the most in this phase (Figure 10C).

**Figure 10.**
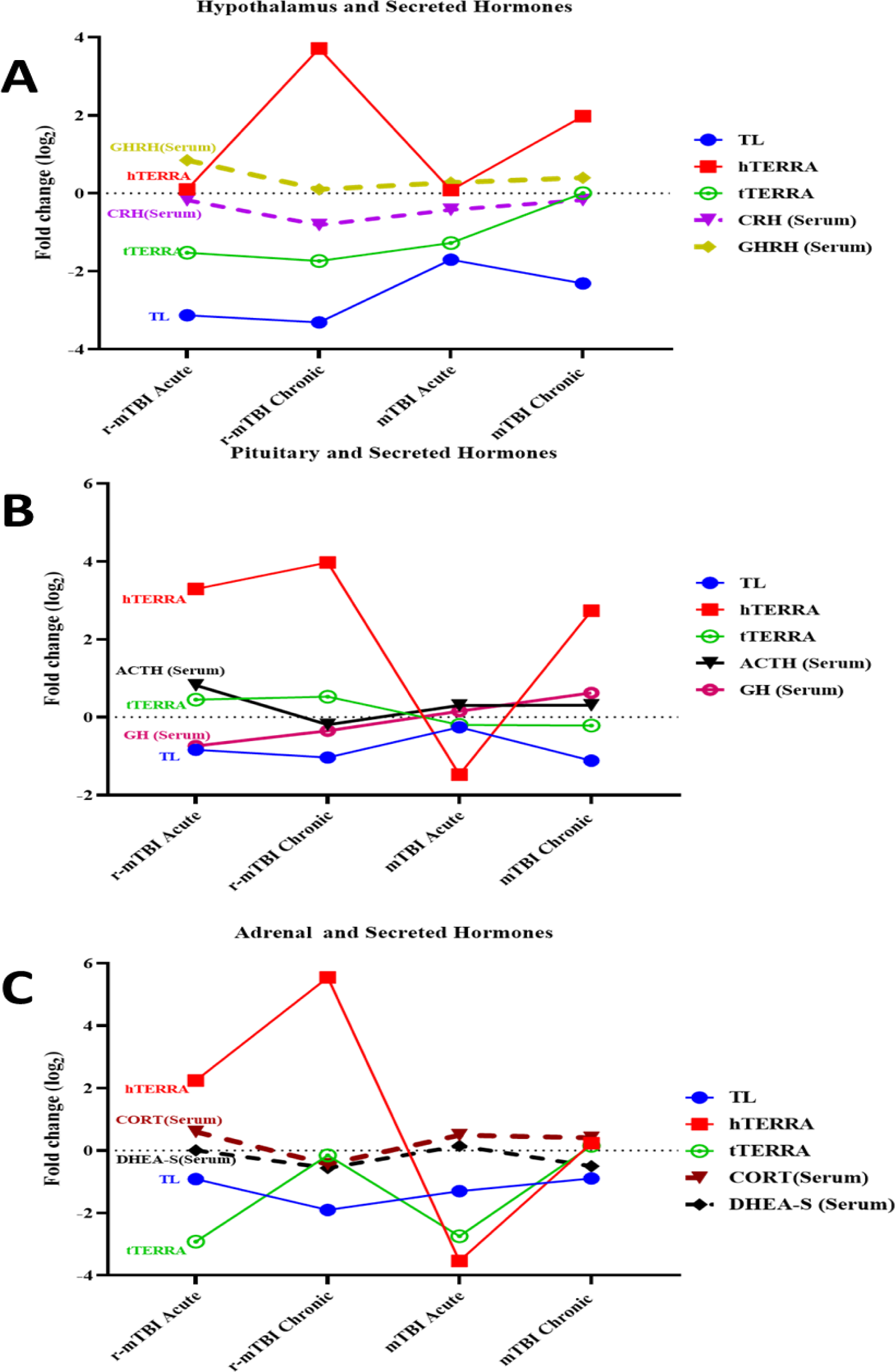
Representation of the relationship between hTERRA, fTERRA, and TL changes in hypothalamic-pituitary and adrenals and serum hormone levels in the mTBI and r-mTBI groups (Values calculated using control values. Baseline=0). Correlations between telomere length, fTERRA, and hTERRA levels, and the levels of hormones released from the hypothalamus, pituitary, and adrenal glands of mTBI mouse groups. When hypothalamic telomere length, fTERRA, and hTERRA levels were compared with CRH and GHRH hormone levels, r-mTBI chronic group had the most shortening of TL and the highest increase hTERRA levels in the hypothalamus, and lowest serum concentrations of CRH and GHRH compared to other groups (Figure 10A). When telomere length, fTERRA and hTERRA levels in the pituitary are compared with GH and ACTH levels; While hTERRA was at its lowest level in the mTBI acute group, ACTH and GH levels were found to be close to the control group. In r-mTBI chronic group demonstrated the shortest telomere length and highest hTERRA levels, GH and ACTH serum levels were decreased (Figure 10B). In the adrenal gland, r-mTBI chronic group had the highest hTERRA levels and, CORT and DHEAS levels decreased, and telomere length shortened the most in this phase (Figure 10C).

The gene transcript levels, telomere length, and free and hybrid TERRA levels obtained from the hypothalamus, pituitary, and adrenal tissues, along with, as well as blood hormone levels, have been summarized in Figure 11.

**Figure 11.**
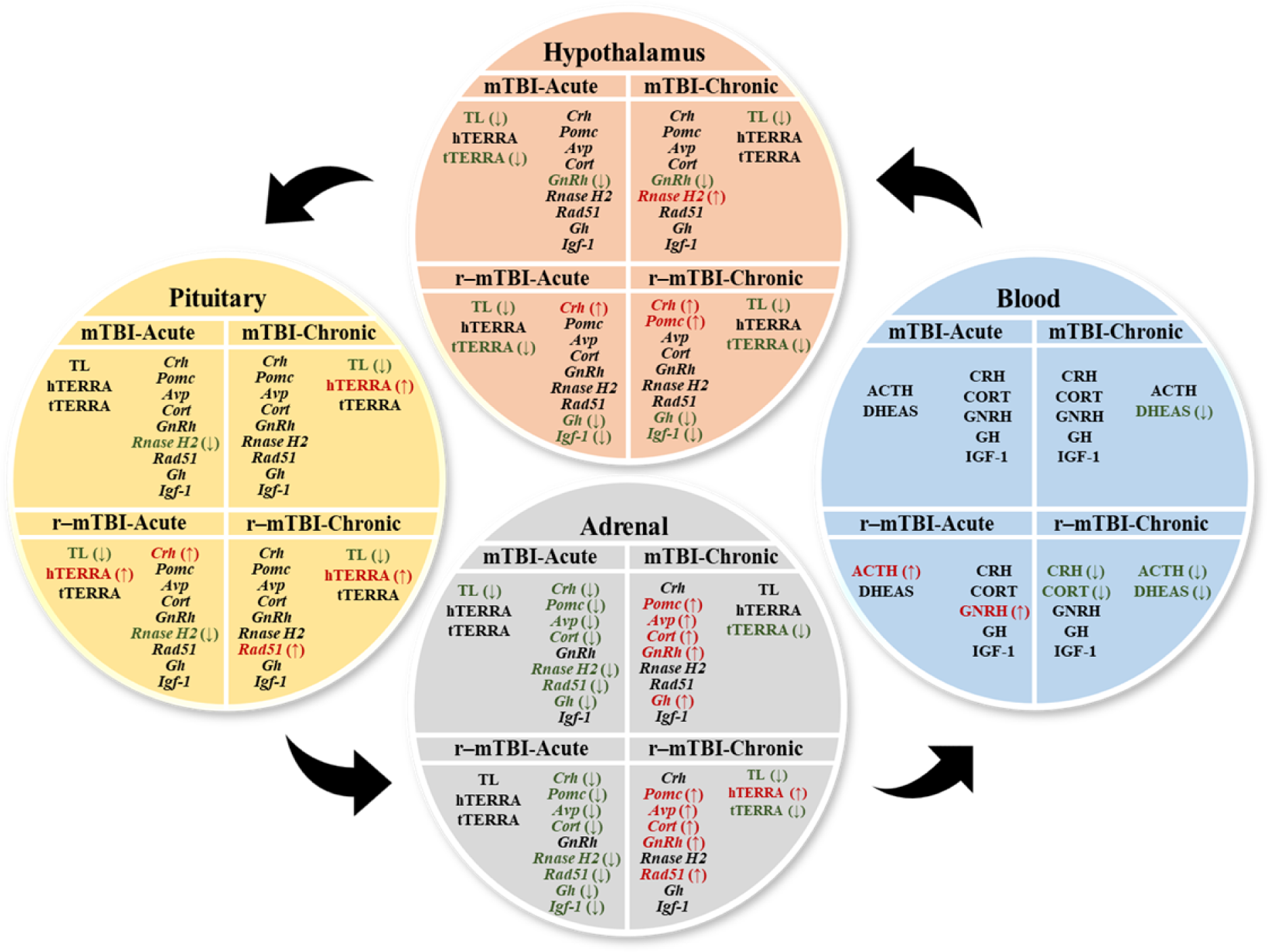
Summary of transcriptional analyses of the hypothalamus, pituitary, and adrenal glands after mTBI and r-mTBI, as well as hormones in the blood (**Green** is up-regulation, **red** is down-regulation).

## Discussion

Traumatic brain injury (TBI) represents a significant public health problem worldwide. Although most people with mild to moderate TBI recover relatively quickly, a substantial subset of individuals experience some debilitating symptoms during the chronic phase. Due to the response to physiological and psychological stressors after TBI, individuals with TBI may suffer from poor stress tolerance, impairments in the ability to appraise stressors, and poor initiation (and termination) of neuroendocrine responses to stress, which can exacerbate TBI-induced stressor dysfunction, especially in the chronic phase. The effects of severe TBI are not difficult to predict. Yet, today we know very little about the effects of mild TBI, to which young children are frequently exposed, especially the impact of repetitive TBI (Tanriverdi *et al*, 2010). Our previous studies showed that pituitary insufficiency occurred after repeated TBI in boxers and kickboxers (Tanriverdi *et al*, 2007; Kelestimur *et al*, 2004). However, at present, the reason why pituitary deficiency occurs and the response mechanisms are not yet known.

In this study, specifically, whether telomere regulation changes in the acute and chronic phases after single or repeated exposure to mTBI, its modulatory role in the acute and chronic phase after TBI, and we will discuss their relationship with HPA, HPG, and GH-IGF-1 axes.

After creating a single and repeated mTBI model in mice, in acute and chronic phase; we first determined the levels of telomere length (TL), total and TERRA transcripts (total and hybridized with DNA), *Rad51*, and *RNase* HII. Then, we also determined the transcript and serum hormone levels associated with the HPA, HPG, and GH axes of the mice. The main objective of this study was to elucidate the mechanism of emergence of diseases associated with TBI and occurring in the chronic phase. There is evidence that cellular events occurring in the hypothalamus, pituitary, and adrenal glands after head injury contribute to the emergence of neurological, hormonal, and psychological problems that occur after TBI.

When we first examined the effect of single and repeated head trauma on the TLs of the hypothalamus, pituitary gland and adrenal glands, we found that the TLs were significantly shortened in the hypothalamus in all trauma groups compared to the control group. The most important point here was that a single mTBI was enough to shorten the TL by ¼ in the hypothalamus. However, the maximum shortening is in the chronic phase of r-mTBI in the hypothalamus. At the pituitary level, it was demonstrated that the highest rate of shortening was observed in the chronic groups compared to the control group. However, the first effect comes from the adrenal, which is consistent with our previous study (Taheri *et al*, 2022a), that is, the highest rate of telomere shortening occurred in the acute phase after a single head trauma.

Indeed, in our previous study, we showed that apoptosis increased during the acute phase at the level of the adrenal glands, which are not tissues directly targeted by head trauma, but that apoptosis increased at the level of the adrenal glands and pituitary gland during the chronic phase (Taheri *et al*, 2022a).

Previous studies have shown that excitotoxic reactions following mTBI can cause specific damage to the hippocampus and hypothalamus, and this damage is associated with cognitive impairment, hormonal dysfunction, and disruption of circadian rhythm observed in clinical and experimental contexts (Kuzminskaite *et al*, 2021). Additionally, adolescent athletes exposed to repetitive mTBI have been shown to have deficits in verbal fluency, memory, cognitive processing, and executive functions (Sezgin Caglar *et al*, 2019). Furthermore, fatigue, insomnia, fragmented sleep, daytime sleepiness, and irregular sleep-wake cycles have also been widely documented in patients with mTBI, and these circadian rhythm disturbances have been associated with hypothalamic lesions (Tanriverdi & Kelestimur, 2015). One of the most important functions of the hypothalamus is homeostasis, it helps ensure the stability of the body. Damage to the hypothalamus after mTBI has been suggested to occur through significant axonal damage or damage to cell bodies, leading to atrophy or apoptosis (Tanriverdi *et al*, 2015, 2006). As we know, telomere length is the biological clock that determines the lifespan of the cell. Cells with short telomeres typically undergo apoptosis (Yamakawa *et al*, 2019). Indeed, the hypothalamus, the pituitary gland and the adrenal glands are closely interacting, and the hormonal response created by the HPA axis against any stress factor is essential for maintaining homeostasis in the body. As a result of any damage, transcriptional and hormonal communication between the hypothalamus, pituitary gland, and adrenal glands is impaired or lost, which can alter the response to many stressors (Taheri *et al*, 2022a).

R-loops, three-stranded DNA/RNA hybrid structures formed during transcription, play a crucial role in transcription and genome integrity. We attempted to reveal how TERRA, the lncRNA transcribed from subtelomeric regions and forming R-loop structures in telomeric regions, varies following trauma in the hypothalamus, pituitary, and the adrenal glands and its relationship with TL. At this stage, the correct regulation of telomeres of the hypothalamus, pituitary gland, and adrenal glands, which play an important role in maintaining the body’s homeostasis with the secretion of various hormones, is very important for the functioning of cells. Recent data reveal the role of persistent or unprogrammed R-loops in telomere regions in supporting genome and telomere instability and their links to human diseases (Gambelli *et al*, 2023).

We determined the levels of free TERRA (fTERRA) and hybrid TERRA (hTERRA) that form R-loop structures after single and repetitive TBI. We propose the relationship between altered telomere regulation of the hypothalamus, pituitary and adrenal glands after head trauma with the HPA, HPG and GH axes. When fTERRA transcript levels in the hypothalamus were compared between groups, they were decreased in all groups, except the mTBI-chronic group, compared to the control group. After a single TBI, the transcription level decreased during the acute period but returned to normal during the chronic period, but repeated TBI could not be tolerated. Although there was no difference between groups regarding fTERRA transcript levels in the pituitary, fTERRA transcript levels decreased significantly in the acute phases in the adrenals and, the transcript levels fTERRA returned to normal during the chronic period. We found that although hTERRA transcript levels usually low, hTERRA levels are markedly increased in the hypothalamus, pituitary and adrenal glands in the chronic phase following head trauma. Furthermore, hTERRA levels were significantly increased in the rmTBI-acute group in the pituitary compared to the mTBI-acute and control groups. Typically, R-loop formations remain in telomere regions for a very short time during DNA synthesis. However, this period is extended in case of DNA damage or replication stress (Gong & Liu, 2023). Our data indicate that with shortened telomeres, the time of R-loop formation is prolonged due to replication stress that occurs particularly in the pituitary and adrenal glands in the chronic phase after head trauma. It plays an important role in R-loop formation in the telomeres of RNaseHII and Rad51. We found that *Rad51* transcript levels and hTERRA levels increased during the chronic phase of the pituitary after r-mTBI. *RNAseHII* transcript levels were also decreased in the adrenal, particularly in acute phases.

It is well-known that the cortisol response to any stressor is essential for the body’s homeostasis and is mediated by the activity of the HPA axis. According to the literature, when the body is confronted with a stressor, the release of CRH by the hypothalamus triggers the release of ACTH and AVP by the pituitary gland. Finally, ACTH triggers the release of cortisol from the adrenals. After a while, an increase in cortisol hormone levels activates the feedback mechanism in the hypothalamus and pituitary gland, thereby suppressing the HPA axis. Along with all this, activation of the HPA axis also affects the HPG and GH-IGF-1 axis. Typically, the HPG and GH-IGF-1 axes are suppressed with the activation of the HPA axis. Although the literature (Wikgren *et al*, 2012) suggests that increased cortisol levels trigger cellular aging, some publications (Savolainen *et al*, 2015) do not support this view. In light of all this previous knowledge, we aimed to reveal the relationship between telomere regulation and the level of transcripts and hormones related to the HPA, HPG and GH-IGF-1 axes after TBI.

When HPA axis-related transcript and hormone levels were compared between groups after mTBI; it was determined that the transcription level of *Crh* increased in the chronic phase after repeated head trauma in the hypothalamus, increased in the acute phase after repetitive mTBI in the pituitary, and decreased in the acute phases in the adrenal glands. Serum CRH levels were also observed to decrease during the chronic phase after repeated head trauma compared to the control group. However, it is interesting to note that in the chronic phase after repetitive mTBI, increased *Crh* transcript levels in the hypothalamus was observed despite the decrease in serum CRH levels. We propose that this is due to the suppression of the body’s CRH response to repetitive stress as a defense mechanism. On the other hand, it was determined that the *Pomc* transcript levels decreased in the hypothalamus in the r-mTBI chronic phase, and in the adrenal gland in acute phases, while it increased in the chronic phases. Although *Pomc* transcript levels decreased in both acute and chronic phases in the pituitary after repeated head trauma, this was not statistically significant. However, serum ACTH levels decreased during the chronic phase after repeated mTBI. Cort transcript levels were not different between groups in the hypothalamus and pituitary but decreased in the adrenals glands in acute phases and increased in chronic stages. In contrast, levels of the cortisol decreased during the chronic phase after repeated mTBI, as did the hormones CRH and ACTH.

As a result of our study, it was determined that cortisol levels increased in the mTBI-acute, mTBI-chronic and r-mTBI acute groups compared to the repeated r-mTBI-chronic phase and the control group. However, TL was shortened after mTBI in all tissues. The data we have obtained so far supports the literature (Wikgren *et al*, 2012; Révész *et al*, 2014). However, the decrease in cortisol levels in the chronic phase after repeated TBI may be related to the fact that hormone-secreting cells may lost their function because maximal telomere shortening occurs in the chronic phase after repeated mTBI in the adrenal glands. In addition, the decrease in CRH and ACTH levels in the chronic phase after r-mTBI, the most shortening of TL in the hypothalamus, pituitary, and adrenals glands were found in the r-mTBI chronic group compared to the control group. Our previous study showed that the first response after TBI comes from the adrenal glands and that markers of apoptosis increase in the adrenals in the acute phase and in the pituitary in the chronic phase (Taheri *et al*, 2022a). Furthermore, the fact that the transcript levels of *Crh, Pomc, Avp,* and *Cort* showed the same pattern in all adrenal groups after single and repeated mTBI showed a clear transcriptional response of the adrenal glands to mTBI rather than the hypothalamus and the pituitary gland.

Studies of the hypothalamic–pituitary–gonadal (HPG) axis describe the regulation of reproductive hormones by the hypothalamus, anterior pituitary gland, and gonads. Through their secretion of essential hormones, the HPG axis profoundly affects development, reproduction, and aging. Systemic inflammation can have a profound effect on the HPG axis. In the hypothalamus, gonadotrophin-releasing hormone (GnRH) is secreted in a pulsatile manner into the pituitary-portal circulation. GnRH acts via the GnRH receptor (GnRHR) on pituitary gonadotrophs to stimulate the synthesis and secretion of luteinizing hormone (LH) and follicle-stimulating hormone (FSH), which then stimulate gonadal steroidogenesis and regulate the gametogenesis. Pathological conditions that cause the secretion of proinflammatory cytokines, such as TBI, impact the activity of principal neurohormonal axes, including the HPA and HPG axes. Additionally, neuropeptide hormones, which regulate HPA axis activity, affect HPG axis function by regulating GnRH secretion by hypothalamic neurons. Glucocorticosteroids, the end products of the HPA axis, have been shown to inhibit GnRH release in the pituitary portal system of the hypothalamic median eminence. However, recent studies suggest that the mechanism underlying the influence of the HPA axis on GnRH neuronal activity is more complex than previously thought (Gołyszny *et al*, 2022).

DHEAS plays an essential role in producing the male sex hormone testosterone and the female sex hormone estrogen. DHEA is produced centrally as a neurosteroid and peripherally by the adrenal cortex and gonads. DHEA also fluctuates in response to social stressors. Unlike other steroid hormones with specific receptors, DHEA binds to several receptors, including the estrogen receptor. DHEA acts as a prohormone for estrogen and testosterone and can broadly affect physiology through its conversion to other sex steroids and metabolites. In one study, they determined brain and plasma levels of neuroactive steroids in young female mice 24 hours, 72 hours and 2 weeks after TBI using the head injury weight-drop model and analyzed whether levels of neuroactive steroids in the brain and plasma after TBI correlated with neurological scores of the animals. They showed that TBI caused neurological deficits detectable 24 and 72 hours after brain injury, which disappeared 2 weeks after injury. They also showed that brain levels of progesterone, tetrahydroprogesterone (THP), isopregnanolone, and 17β-estradiol decreased 24 hours, 72 hours, and 2 weeks after TBI, while levels of DHEA and testosterone decreased temporarily 24 hours after injury. As a result of their analysis, they determined that local levels of brain progesterone and DHEA had a positive correlation with neurological recovery. Consistent with the data obtained, they showed that TBI modifies the levels of neuroactive steroids in the brain independently of plasma levels, and they suggested that increasing the levels of progesterone and DHEA in the brain could promote neurological recovery after TBI (Lopez-Rodriguez *et al*, 2015). Therefore, at the transcriptional level, this might actually be involved in the regulating of the HPG axis response that is not reflected in the periphery after TBI, and changes in GnRH transcript levels, especially in the hypothalamus and adrenal glands, could be a sign of this.

In our study, serum DHEAS levels were determined as well as *GnRH* transcription levels in the hypothalamus, pituitary and adrenal glands after single and repetitive TBI. While *GnRH* transcription levels in the hypothalamus decreased in all groups compared to the control group, the highest decrease occurred in the chronic phase after a single head injury. Although there was no difference between groups in the pituitary gland, it was found that the most significant change in *GnRH* transcript levels occurred in the adrenal glands and that the levels of *GnRH* transcription were significantly increased in the chronic phase compared to the control and acute groups. Transcription levels of *GnRH* in the hypothalamus decreased in all groups compared to the control group. It was found that the greatest decrease occurred in the chronic phase after a single mTBI. However, it was remarkable that DHEAS levels decreased during the chronic phase after repetitive head injury among the groups after mTBI. However, there was a significant increase in hTERRA levels with maximal telomere shortening in the hypothalamus of the chronic group with repetitive mTBI in the hypothalamus. This may explain the decreased DHEAS levels. The study showed that mice with growth hormone deficiency (IGHD) lived longer than their normal siblings. They have also been shown to age more slowly and increase their longevity. The negative association between GH signaling and longevity apparently extends to other mammalian species, including humans. Proportional short stature, doll-like facial features, high-pitched voices, and central obesity characterize people with IGHD. Reach puberty late but are fertile and generally healthy. Additionally, these IGHD people are partially protected against cancer and some of the common effects of aging. One case of IGHD lived to be 103 years old. Researchers believed that the secret to these individuals’ longevity might be due to a permanent reduction in circulating IGF-1, low levels of IGF-2, and decreased GH secretion (Aguiar-Oliveira & Bartke, 2019).

Our study also determined the transcripts and hormones associated with the GH-IGF-1 axis in mice in the acute and chronic phases after single and repetitive mTBI. It was found that after repeated head trauma, *Gh* transcript levels in the hypothalamus are excessively decreased in the acute phase and especially in the chronic phase. Although there was no significant difference between groups at the pituitary level, the decrease in *Gh* transcript levels in the acute phases after single and repeated mTBI to the adrenal glands is remarkable. There was no significant difference in *Igf1* transcript levels in the pituitary and adrenal glands, *Igf-1* transcript levels decreased in the acute and chronic phases after rmTBI in the hypothalamus. However, changes in transcription levels in the hypothalamus were not reflected in the periphery. There was no significant difference between the serum levels of GH and IGF-1 of the groups, but it was found that serum GHRH levels increased in the acute phase after repetitive mTBI. GHRH is a hormone produced in the hypothalamus. The primary role of GHRH is to stimulate the pituitary gland to produce and release GH into the bloodstream. Therefore, increased GHRH levels may have been in response to a decrease in serum lGH levels. Additionally, a study examined whether a single or repetitive TBI (after four repetitive injuries) led to disruption of the GH/IGF-1 axis in mice. Basal levels of circulating GH and IGF-1 were measured at baseline at 24 hours, 72 hours, 1 week, and 1 week after TBI. The results revealed that r-mTBI exhibited a significant acute and chronic reduction in circulating GH and IGF-1 compared to sham and simple-mTBI animals (Greco *et al*, 2013). We suggest that the reason why we could not find a difference between serum levels of GH and IGF-1 in our study, unlike the previous research, is related to the severity of TBI.

Our data indicate that serum levels of the HPA, HPG, and GH-IGF-1 axis-associated hormones decreased significantly during the chronic phase after repetitive mTBI, as expected. These results prove pituitary insufficiency occurs in the chronic phase after head trauma. However, when changes in the transcription levels of these hormones are assessed locally, the local increase in the transcription levels of the *Crh* and *Pomc* genes, in particular, suggests a defensive response through transcriptional alteration against mTBI, despite the decreased rate in the serum in chronic phase. Additionally, changes in the transcription levels of TERRA molecules, responsible for the regulation of telomere length and R-loop formation, as well as their correlation with hormonal deficits after repetitive mTBI head trauma, were observed. Especially after repeated mTBI, the maximal shortening of telomeres and the maximal increase in hTERRA level in the chronic phase suggest a possible disorder of genome stabilization and loss of cellular function in tissues after a TBI in the hypothalamus, pituitary gland, and adrenal glands. The data we acquired will significantly contribute to the elucidation of the mechanism of TBI-induced diseases, especially pituitary insufficiency.

Our data indicated that single-mTBI and r-mTBI disrupt telomere regulation and telomeres shortening via TERRA in cells of the hypothalamus, pituitary, and adrenal glands. Telomere shortening and dysregulation of TERRA in the hypothalamus, pituitary, and adrenal glands results in impaired genome integrity and cellular function. As a result, hormonal regulation of the HPA, HPG, and GH-IGF-1 axes is disrupted in the chronic phase after mTBI. Impaired and shortened telomere regulation due to TBI may underlie TBI-related diseases, especially pituitary insufficiency.

Finally, these studies will contribute to the development of new treatment strategies as well as diagnosis and monitoring methods after head trauma.

## Author Contributions

F.K., S.T., M.R., Z.K., F.T., K.U., and Z.Y. conceived and designed the experiments and the primary research hypothesis. Z.Y., E.M., M.M., and B.E. designed and applied the TBI model and collected tissues. Z.Y. and B.E. performed the experiments. Z.Y. and M.G.Ö. performed the statistical analysis and data visualization. F.K., S.T., M.R., Z.K., F.T., and K.U. analyzed the draft manuscript and corrected its style and content. F.K., S.T., M.R., Z.K., F.T., and K.U. supervised all steps of this study, including the experiments. All the aforementioned authors fully contributed to the writing, reading and approval of the final version of this manuscript.

## Data Availability

Data can be shared by contacting the corresponding author upon the request of the researchers.

## References

Aguiar-Oliveira MH & Bartke A (2019) Growth Hormone Deficiency: Health and Longevity. Endocr Rev 40: 575–601

Akkar I, Karaca Z, Taheri S, Unluhizarci K, Hacioglu A & Kelestimur F (2022) The stimulatory effects of glucagon on cortisol and GH secretion occur independently from FGF-21. Endocrine 75: 211–218

Bazaz MR, Balasubramanian R, Monroy-Jaramillo N & Dandekar MP (2021) Linking the Triad of Telomere Length, Inflammation, and Gut Dysbiosis in the Manifestation of Depression. ACS Chem Neurosci 12: 3516–3526

Dalal M & Khanna-Chopra R (2001) Differential response of antioxidant enzymes in leaves of necrotic wheat hybrids and their parents. Physiol Plant 111: 297–304

De Rosa M & Opresko PL (2023) Translating the telomeres. Trends Genet TIG 39: 593–595

Feretzaki M, Pospisilova M, Valador Fernandes R, Lunardi T, Krejci L & Lingner J (2020) RAD51-dependent recruitment of TERRA lncRNA to telomeres through R-loops. Nature 587: 303–308

Gambelli A, Ferrando A, Boncristiani C & Schoeftner S (2023) Regulation and function of R-loops at repetitive elements. Biochimie 214: 141–155

Gołyszny M, Obuchowicz E & Zieliński M (2022) Neuropeptides as regulators of the hypothalamus-pituitary-gonadal (HPG) axis activity and their putative roles in stress-induced fertility disorders. Neuropeptides 91: 102216

Gong Y & Liu Y (2023) R-Loops at Chromosome Ends: From Formation, Regulation, and Cellular Consequence. Cancers 15: 2178

Graf M, Bonetti D, Lockhart A, Serhal K, Kellner V, Maicher A, Jolivet P, Teixeira MT & Luke B (2017) Telomere Length Determines TERRA and R-Loop Regulation through the Cell Cycle. Cell 170: 72–85.e14

Greco T, Hovda D & Prins M (2013) The Effects of Repeat Traumatic Brain Injury on the Pituitary in Adolescent Rats. J Neurotrauma 30: 1983–1990

Isken O & Maquat LE (2009) Telomeric RNAs as a novel player in telomeric integrity. F1000 Biol Rep 1: 90

Kelestimur F, Tanriverdi F, Atmaca H, Unluhizarci K, Selcuklu A & Casanueva FF (2004) Boxing as a sport activity associated with isolated GH deficiency. J Endocrinol Invest 27: RC28-32

Kuzminskaite E, Penninx BWJH, van Harmelen A-L, Elzinga BM, Hovens JGFM & Vinkers CH (2021) Childhood Trauma in Adult Depressive and Anxiety Disorders: An Integrated Review on Psychological and Biological Mechanisms in the NESDA Cohort. J Affect Disord 283: 179–191

Lipps HJ & Rhodes D (2009) G-quadruplex structures: in vivo evidence and function. Trends Cell Biol 19: 414– 422

Lopez-Rodriguez AB, Acaz-Fonseca E, Giatti S, Caruso D, Viveros M-P, Melcangi RC & Garcia-Segura LM (2015) Correlation of brain levels of progesterone and dehydroepiandrosterone with neurological recovery after traumatic brain injury in female mice. Psychoneuroendocrinology 56: 1–11

Marchlewska-Koj A, Kruczek M & Toch E (1983) Suppression of estrous cycle of female mice by ovariectomized females. Horm Behav 17: 233–236

Marmarou A, Foda MA, van den Brink W, Campbell J, Kita H & Demetriadou K (1994) A new model of diffuse brain injury in rats. Part I: Pathophysiology and biomechanics. J Neurosurg 80: 291–300

Mehmetbeyoglu E, Kianmehr L, Borlu M, Yilmaz Z, Basar Kılıc S, Rajabi-Maham H, Taheri S & Rassoulzadegan M (2022) Decrease in RNase HII and Accumulation of lncRNAs/DNA Hybrids: A Causal Implication in Psoriasis? Biomolecules 12: 368

O’Callaghan NJ & Fenech M (2011) A quantitative PCR method for measuring absolute telomere length. Biol Proced Online 13: 3

Rassoulzadegan M, Sharifi-Zarchi A & Kianmehr L (2021) DNA-RNA Hybrid (R-Loop): From a Unified Picture of the Mammalian Telomere to the Genome-Wide Profile. Cells 10: 1556

Révész D, Verhoeven JE, Milaneschi Y, de Geus EJCN, Wolkowitz OM & Penninx BWJH (2014) Dysregulated physiological stress systems and accelerated cellular aging. Neurobiol Aging 35: 1422–1430

Savolainen K, Eriksson JG, Kajantie E, Lahti J & Räikkönen K (2015) Telomere length and hypothalamic-pituitary-adrenal axis response to stress in elderly adults. Psychoneuroendocrinology 53: 179–184

Sezgin Caglar A, Tanriverdi F, Karaca Z, Unluhizarci K & Kelestimur F (2019) Sports-Related Repetitive Traumatic Brain Injury: A Novel Cause of Pituitary Dysfunction. J Neurotrauma 36: 1195–1202

Taheri S, Karaca Z, Mehmetbeyoglu E, Hamurcu Z, Yilmaz Z, Dal F, Çınar V, Ulutabanca H, Tanriverdi F, Unluhizarci K, et al (2022a) The Role of Apoptosis and Autophagy in the Hypothalamic-Pituitary-Adrenal (HPA) Axis after Traumatic Brain Injury (TBI). Int J Mol Sci 23: 15699

Taheri S, Karaca Z, Rassoulzadegan M, Mehmetbeyoglu E, Zararsiz G, Sener EF, Bayram KK, Tufan E, Sahin MC, Marasli MK, et al (2022b) The Characterization of Sex Differences in Hypoglycemia-Induced Activation of HPA Axis on the Transcriptomic Level. Cell Mol Neurobiol 42: 1523–1542

Taheri S, Tanriverdi F, Zararsiz G, Elbuken G, Ulutabanca H, Karaca Z, Selcuklu A, Unluhizarci K, Tanriverdi K & Kelestimur F (2016) Circulating MicroRNAs as Potential Biomarkers for Traumatic Brain Injury-Induced Hypopituitarism. J Neurotrauma 33: 1818–1825

Tanriverdi F & Kelestimur F (2015) Pituitary dysfunction following traumatic brain injury: clinical perspectives. Neuropsychiatr Dis Treat 11: 1835–1843

Tanriverdi F, Schneider HJ, Aimaretti G, Masel BE, Casanueva FF & Kelestimur F (2015) Pituitary dysfunction after traumatic brain injury: a clinical and pathophysiological approach. Endocr Rev 36: 305–342

Tanriverdi F, Senyurek H, Unluhizarci K, Selcuklu A, Casanueva FF & Kelestimur F (2006) High risk of hypopituitarism after traumatic brain injury: a prospective investigation of anterior pituitary function in the acute phase and 12 months after trauma. J Clin Endocrinol Metab 91: 2105–2111

Tanriverdi F, Unluhizarci K, Coksevim B, Selcuklu A, Casanueva FF & Kelestimur F (2007) Kickboxing sport as a new cause of traumatic brain injury-mediated hypopituitarism. Clin Endocrinol (Oxf) 66: 360–366

Tanriverdi F, Unluhizarci K & Kelestimur F (2010) Pituitary function in subjects with mild traumatic brain injury: a review of literature and proposal of a screening strategy. Pituitary 13: 146–153

Tufan E, Taheri S, Karaca Z, Mehmetbeyoglu E, Yilmaz Sukranli Z, Korkmaz Bayram K, Ulutabanca H, Tanrıverdi F, Unluhizarci K, Rassoulzadegan M, et al (2023) Alterations in Serum miR-126-3p Levels over Time: A Marker of Pituitary Insufficiency following Head Trauma. Neuroendocrinology: 1–16

Tweedie D, Rachmany L, Rubovitch V, Zhang Y, Becker KG, Perez E, Hoffer BJ, Pick CG & Greig NH (2013) Changes in mouse cognition and hippocampal gene expression observed in a mild physical-and blast-traumatic brain injury. Neurobiol Dis 54: 1–11

Wikgren M, Maripuu M, Karlsson T, Nordfjäll K, Bergdahl J, Hultdin J, Del-Favero J, Roos G, Nilsson L-G, Adolfsson R, et al (2012) Short telomeres in depression and the general population are associated with a hypocortisolemic state. Biol Psychiatry 71: 294–300

Yamakawa GR, Weerawardhena H, Eyolfson E, Griep Y, Antle MC & Mychasiuk R (2019) Investigating the Role of the Hypothalamus in Outcomes to Repetitive Mild Traumatic Brain Injury: Neonatal Monosodium Glutamate Does Not Exacerbate Deficits. Neuroscience 413: 264–278

